# The Statistical Foundation of Colour Space

**DOI:** 10.1101/849984

**Authors:** Lucas Wilkins

**Affiliations:** University of Oxford

**Keywords:** Fisher Information, Information Geometry, Colour Vision, Bee Vision

## Abstract

The idea of a colour space where distance corresponds to discriminability has been fundamental to colour vision research since the 19^th^ century. Despite their long-standing success there is a contradiction between the geometric framework that is typically used in these spaces (a particular application of Riemannian geometry) and a view of the transduction of sensory information as the result of a stochastic process. When this is made explicit, a subtly different approach is suggested which turns out to provide a general, and more complete framework for colour space. It is argued further that not only is a contradiction avoided, but that this framework is both intuitive and of real practical value, in particular for researchers interested in the visual behaviour and ecology of animals.

## 1 Introduction

Spatial thinking about colour has a long history. Written accounts of spatial arrangements of colour, such as ‘colour wheels’, date back at least to the 13^th^ century[11], and there are diagrams which could be called colour spaces that survive from 1700s. Early representations of relationships between colours, such as how pigments are affected by mixing, were based on simple geometric shapes: circles, prisms, cones, spheres etc. and were precursors to modern colour spaces, which have tended to grow in detail and sophistication, gradually incorporating of our increasing understanding of colour, and being tailored to different applications.

A significant change in the conception colour spaces occurred in the period around the start of the 20^th^ century; arising as a combination of new ideas in geometry and the psychophysics of the mid 1800s. Fundamental to this shift was the work of Gustav Fechner which linked physical measures of magnitude with psychological scales describing sensation[5]. This work embodied the principle that a subjects ability to distinguish stimuli corresponds to their relative position on a perceptual scale (their sensation). Two pairs of stimuli that are equally distinguishable are equally distant in the perceptual scale. This meant that a geometric arrangement of colours could be made, with distance corresponding to the ease with which they can be distinguished. Colour diagrams had evolved into a kind of “perceptual space”.

This idea found its formal expression in the Riemann’s formulation of the geometry of curved surfaces and spaces; and it was Helmholtz who first took these new ideas in geometry to make a trichromatic space based on Fechner’s principle[13]. To Helmholtz value of this geometry lied in the intuitiveness of spatial arrangement and the flexibility of a space that can be warped and stretched, allowing the distance from each colour to its neighbours to be different for each point. This approach is the foundation of modern colour spaces, and is a methodological approach that makes colour theory and its application different from that of other senses.

Powerful though this approach is, I will explain how current formulations of this kind of space clash with another way of thinking about sensation – as a physiological process of information transduction. Such an approach sees sensory systems a cascade of mechanistic (though stochastic) processes that transduce information from the world outside, and a behavioural experiment measuring the response to stimuli can be thought of as observing the result of information passing through a complex, noisy, mass of biological connections. This paper begins by establishing that this view is incompatible with many widely used colour spaces being correctly described by Riemann’s geometry.

This result, whilst concrete, is not a fatal flaw in established colour spaces, after all they are used to great effect. Rather, it suggests an underlying statistical foundation to the notion of colour space in which established colour spaces can be seen as a kind of approximation, and where Riemann’s geometry is compatible with the physiological view above.

But, if all this foundation provided was a means to rescue a particular kind of geometric approach – one which was only ever used for pragmatic reasons in the first place – it perhaps wouldn’t be that valuable. Why should we care about whether a particular geometric model is strictly applicable, after all, why should we assume there is anything innately geometric about colour? The second half of this aims to render this question moot, by demonstrating how this updated view of colour space paper is useful in of itself. I will explain how this foundation gives a more complete and intuitive picture of what a colour space is and how it relates to physiology, as well as an application to demonstrate its practical value.

I will argue that better correspond to our intuitions of colour spaces, and are really “what we are getting at” when we talk about them, both in terms of the results of colour discrimination experiments and when thinking about the transduction of colour information. Finally, I will show how it provides a rigorous way of obtaining colour spaces based on models of physiological processes, highlighting some facets of this kind of model that would be otherwise problematic. Such models have particular value for animal colour vision, where physiological investigation is more ready than psychological investigation.

This paper cuts across a number of different scientific and mathematical domains, and although much of the work I will build upon is well established and uncontroversial, I expect that the average reader will be unfamiliar with at least some of the topics at hand. For this reason I have prioritised motivation over technical detail, and opted for a discursive mode of explanation that emphasises the scientific context of the mathematical results. Those who desire a more formal treatment of the mathematics should direct their attention first to Amari and Nagaoka’s monograph on information geometry[1].

## 2 Background

Our capacity to distinguish between things using our senses rarely have a simple, linear relationship to units we use to describe the physical world. This is as true for colour as it is for any sense, and the modern idea of a colour space as a means of quantifying discriminability can be traced back to the earliest psychophysical laws.

Weber’s law is first and perhaps most well known description of a psychophysical relationship. Here the threshold of discrimination Δ*s* has an inverse relationship with the magnitude of the stimulus:

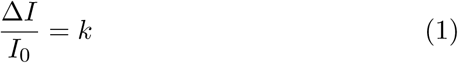

A common interpretation of this is that the physical scale measured in terms of *I*, is logarithmically related to a psychological scale *s* (a scale of sensation) in which the threshold of discrimination is the same for every value. The scale that corresponds to Weber’s law is logarithmic:

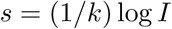

This is known as Fechner’s law, and it is obtained from Weber’s law by solving the following differential equation, obtained as a generalisation equation 1.

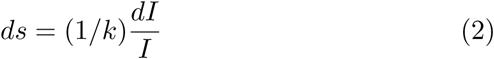

As Fechner’s law is a generalisation of Weber’s law, Weber’s law follows deductively from it. This technique will be important later, so I will describe it a little more explicitly than is strictly necessary at this point.

Weber’s law is obtained from Fechner’s law by a linear approximation of *s* in terms of *I* around some intensity *I*_0_. This amounts to approximating *s* with its tangent line at *I*_0_. We can express this algebraically as:

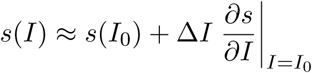

Or, more simply, if we define Δ*s* as *s*(*I*) − *s*(*I*_0_) we can obtain

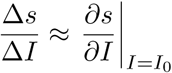

This can then be substituted into 2 to give

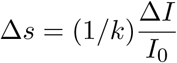

and if we say that the discriminability threshold for *s* is 1 and so set Δ*s* to this value, then we arrive at Weber’s law (eqn. 1).

### 2.1 Fechner’s Law to a Colour Space

Fechner’s law is one dimensional, but colour is multidimensional. To adapt Fechner’s law to the trichromatic vision of humans, Helmholtz used the Pythagorean relationship to combine Fechner’s law acting on different colour channels, using *x, y* and *z* to the intensity of three primary colours that are mixed to form a specific colour.

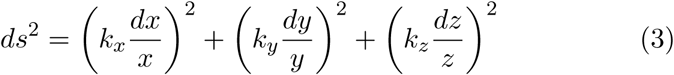

*ds* is known as the *line element*; and it represents the length of an infinitesimal portion of a line^1^. Since Helmholtz, different formulae have been proposed for the line element, the formulae for which all have the same form: a sum of second order infinitesimals, i.e. square terms such as *dx*^2^, or mixed second order terms such as *dy dz*. Each term is weighted by value that depends on one or more coordinates.

We can write the general form of the line element as

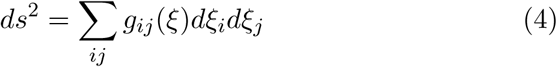

where *ξ* = (*ξ*_1_, *ξ*_2_ … *ξ*_*n*_) is a vector of the parameters specifying a colour, *n* is the number of colour channels. In the case of Helmholtz model of human vision *ξ* is the 3-dimensional vector (*x, y, z*). In modern treatments *ξ* is typically not the intensity of primaries as used by Helmholtz’ but the *quantum catch* – a measure based on the amount of light exciting a particular kind of cone cell defined as:

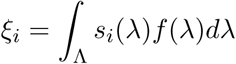

where *s*_*i*_ is the *cone spectral sensitivity function*, also known in the human literature as a cone fundamental, which specifies how effective each wavelength of light is at causing the phototransduction events that lead to a neural signal. The function *f* describes the spectrum of light hitting the eye, and the integral is taken over all visible wavelengths (Λ).

The quantity *dξ* = (*dξ*_1_, *dξ*_2_ … *dξ*_*n*_) is a vector describing an infinitesimal change in the *ξ* parameters, equivalent to (*dx, dy, dz*) in Helmholtz’ model. *g*(*ξ*) is an *n*-by-*n* matrix which is both symmetric (*g*_*ij*_ = *g*_*ji*_) and positive definite. The property of positive definiteness is equivalent to saying *ds*^2^ must always be positive quantity. In the framework of differential geometry, *g*(*ξ*) is known as the metric tensor. The metric tensor is cornerstone of Riemannian geometry, defining it for every point in a space is enough to describe its geometry entirely.

Riemannian geometry is a whole system for describing geometric shape, and the definition of the line element is only one aspect of it. Much of Riemannian geometry is concerned with calculating things like curvature and the shortest path between two distant points, but colour theory does not typically put these tools to much use, being only concerned with the small distances that correspond to the limit of detectability.

There are plenty of possible *n*-by-*n* matrices that can play the role of the metric tensor, and various approaches have been taken in colour theory. In Helmholtz’ models the matrix was obtained from psychophysical laws, but by the mid 20^th^ century, people were deriving the metric tensor more empirically[9, 15]. Thresholds were found by fitting psychometric curves to experimental data, and people were creating spaces by measuring the variability found in tasks involving the matching of colours.

The inverse of a covariance matrix is often explicitly identified as the metric tensor determining a colour space; examples include Wyszecki and Stiles [15] textbook on colour science and the most widely used colour model in animal vision[14]. Moreover, if one takes a decision theoretic view of psychometric curves, then the covariance matrix can be viewed as representing the uncertainty implicit in the decision. For these reasons I have here taken covariance matrices the canonical basis for metric tensor that is used in practice for the construction of colour spaces. I will use this as the starting point for the argument I present here.

We will see in the forthcoming sections that this use of covariance matrices presents a mathematical problem for Reimannian geometry. This problem originates in a third property of the metric tensor which I will discuss below, and is connected to how the metric tensor must responds to changes of coordinate. Before addressing this, it will be useful to quickly review how symmetric, positive-definite matrices can be represented geometrically as ellipses and ellipsoids. This is a widely used and intuitive way of thinking about the kind of quantities that are central to this paper.

#### 2.1.1 Positive Definiteness and Ellipsoids

Like we saw with Fechner’s law a linear approximation can be used to give a finite version of Helmholtz’ line element. If we take consider the threshold to be where Δ*s* = 1 we get the following equation:

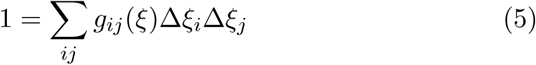

This is the general equation for an ellipse or ellipsoid, the correct term depending on its dimension – ellipses are 2D ellipsoids (I shall just use the term ellipse from here on for the sake of both clarity and brevity). Ellipses are an intuitive way of visualising the metric tensor, and they have been used a lot in human colour vision to this end, the most well known being the MacAdam ellipses for human observers[8].

The fact that equation 5 describes an ellipsoid is entailed by *g* being a positive-definite symmetric matrix. The positive-definiteness of *g* is a requirement of Riemannian geometry and it is what assures that the distance between two points is a positive number.

The positive definiteness of matrices has useful interpretations in terms of ellipses. For instance, if one ellipse lies entirely within another, the difference between the matrices that specify them is positive definite. It is sometimes useful to write this relationship in a way analogous to an inequality: writing

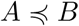

states that *B* − *A* is positive-semidefinite, and so the ellipsoid that *B* describes is contained within the one described by *A*. We can expand this notation to that shown in figure 2.

**Figure 1:**
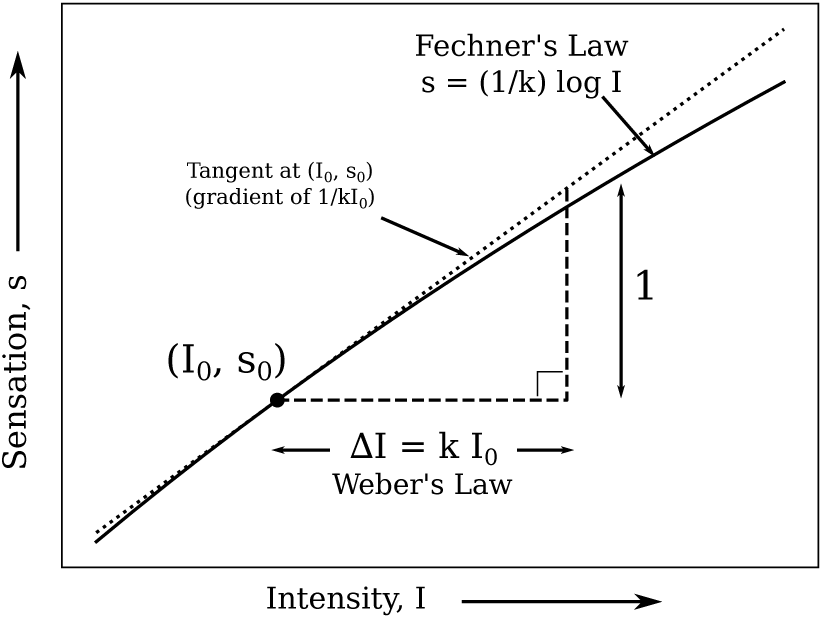
Weber’s law as obtained from linearising Fechner’s law.

**Figure 2:**
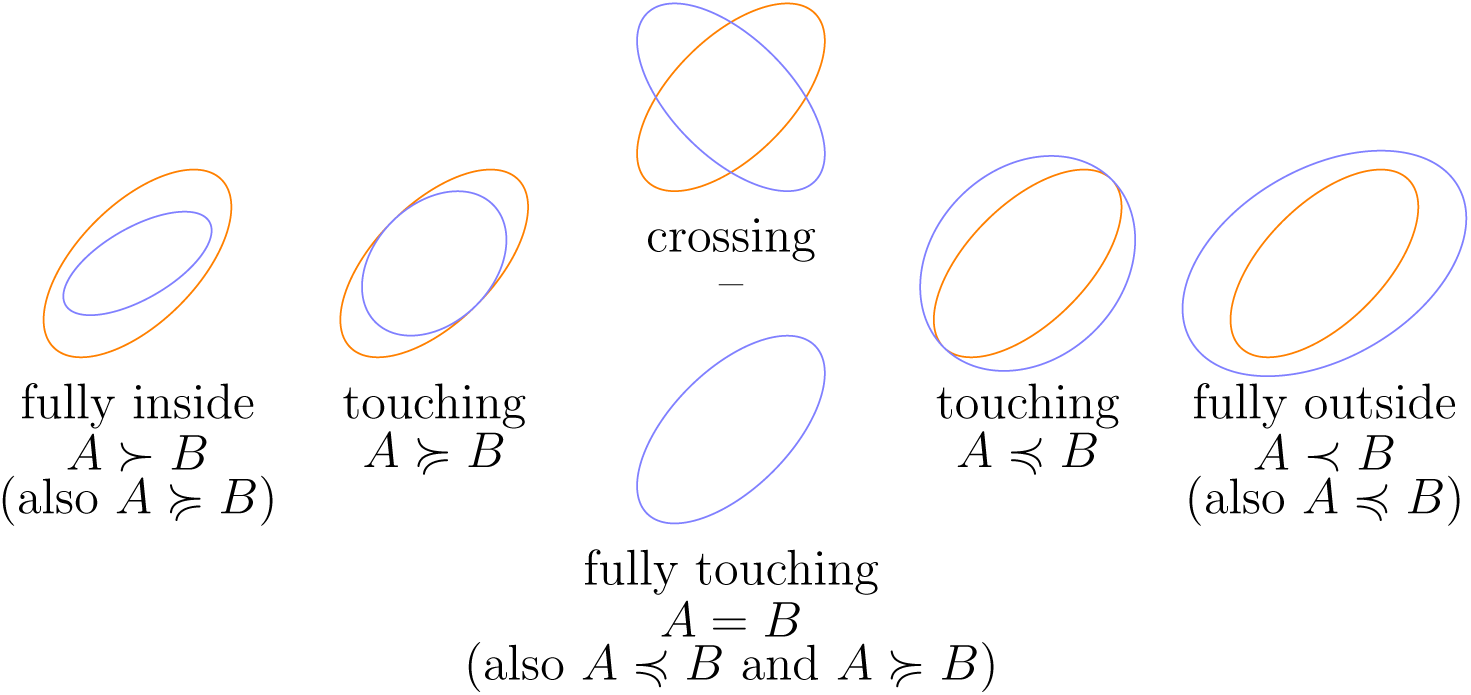
The relative positive-definiteness relation interpreted in terms of two ellipses. Orange ellipses represent a matrix *A* via the equation Σ_*i,j*_ *A*_*ij*_Δ*x*Δ*y* = 1 and blue via the equation Σ_*i,j*_ *B*_*ij*_Δ*x*Δ*y* = 1.

With this notion it is possible to write that a matrix is positive definite by saying that it is positive definite relative to a matrix of zeros:

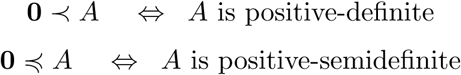

I will make use of this notation in the pages to come.

### 2.2 The Metric Tensor, Covariance and Contravariance

The power of differential geometry lies in the ability to change the coordinate system being used without changing the magnitudes of lines, areas or volumes. Just as the physical distance between London and Paris remains the same whether we represent them using a Mercator map or a gnomic map, the thresholds of discriminability remain the same whether we represent colour using Helmholtz’ *X, Y* and *Z*, the modern CIE coordinates or any other choice; What changes is how we would go about calculate that distance in those coordinates.

In terms of the models of colour we have been discussing this means that the line element *ds* does not depend on the choice of coordinates. Within Riemannian geometry, this is achieved by changing the metric tensor *g* in a manner complementary to the change in coordinates. We can see how this works most easily by considering a one dimensional example with two different coordinate systems. I will distinguish these coordinates by using the letters *ξ* and *ζ*. If we call the metric tensor in the two coordinate systems *g*^Ξ^(*ξ*) and *g*^*Z*^(*ζ*) respectively (the superscript symbols Ξ and *Z* being the uppercase version of the parameter symbols *ξ* and *ζ*), then as *ds* is the same in both, equation 4 gives us

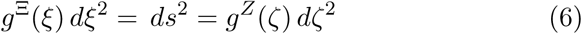

The relationship between *dξ* and *dζ* is given by the chain rule for differentiation

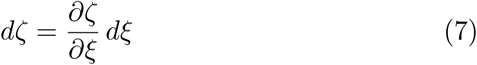

Using this and equation 6 we can see that the metric tensor must change in the ‘opposite’ way if *ds* is to remain constant.

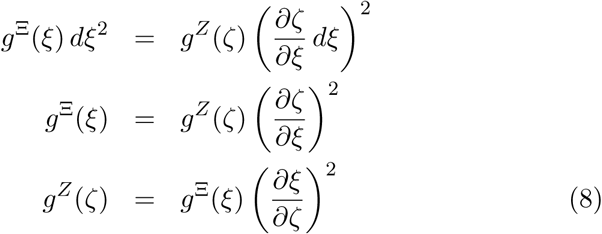

We can see here that to get *g*^*Z*^ from *g*^Ξ^, we need multiply by the inverse derivative of the coordinate transformation twice. One can check this is correct, by substituting it along with 7 into 6, and the partial derivatives of *ξ* with respect to *ζ* will cancel with the derivatives of *ζ* with respect to *ξ*.

In differential geometry there is terminology associated with the way different quantities must be multiplied to keep *ds* the same. A quantity that needs to be multiplied by 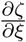 is called a *contravariant tensor*, and one that needs to be multiplied by 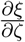 is called a *covariant tensor*. The number of times one must multiply by one of these quantities is called the tensor’s *rank*. The metric tensor is a *rank 2 covariant tensor*. The line element, *ds*, is an *invariant* as it remains the same in all coordinate systems. There are plenty of quantities that are not covariant, invariant or contravariant.

The manner in which the metric tensor and related objects are required to be transformed so as to keep distances the same is central to the first part of my case.

So far I have only discussed a one dimensional example, as formulae for this case are more transparent. However, for multiple dimensions the story is very much the same: instead of multiplying single values, we are multiplying vectors, matrices and so forth. Here is a summary of some the quantities that are used in this paper and how they transform with coordinates:

| Name | Symbol ($\Xi$) | Type | Rank | Transformation (to $Z$) |
| --- | --- | --- | --- | --- |
| Metric Tensor | $g_{ij}^{\Xi}$ | Covariant | 2 | $g_{ab}^Z = \sum_{i,j} \frac{\partial \xi_i}{\partial \zeta_a} \frac{\partial \xi_j}{\partial \zeta_b} g_{ij}^{\Xi}$ |
| Coordinate Derivative | $\frac{\partial}{\partial \xi_i}$ | Covariant | 1 | $\frac{\partial}{\partial \zeta_a} = \sum_i \frac{\partial \xi_i}{\partial \zeta_a} \frac{\partial}{\partial \xi_i}$ |
| Line Element | $ds$ | Invariant | 0 | $ds$ |
| Coordinate Infinitesimal | $d\xi_i$ | Contravariant | 1 | $d\zeta_a = \sum_i \frac{\partial \zeta_a}{\partial \xi_i} d\xi_i$ |
| Inverse Metric Tensor | $(g^{\Xi})_{ij}^{-1}$ | Contravariant | 2 | $(g^Z)_{ab}^{-1} = \sum_{i,j} \frac{\partial \zeta_a}{\partial \xi_i} \frac{\partial \zeta_b}{\partial \xi_j} (g^{\Xi})_{ij}^{-1}$ |

I have not yet talked about the last entry – the inverse of the metric tensor. This will be a more useful quantity for later discussions than the metric tensor itself. This quantity follows a general rule that the inverse of a covariant tensor is contravariant, and vice-versa, with the rank remaining the same.

### 2.3 Perceptual Uniformity

Although the ability to use different coordinate systems is central to differential geometry there are some choices of coordinates that are of special importance to psychophysics. These are those coordinates directly correspond to discriminability. We have already encountered this kind of coordinate system in Fechner’s law.

In this kind of coordinate system, the line element can be written as

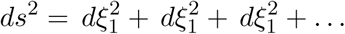

This is where the discrimination ellipses/ellipsoids are circles/spheres of radius 1. For the model given in equation 3 one possible perceptually uniform coordinate system is the coordinates (*ξ*_1_, *ξ*_2_, *ξ*_3_) given by:

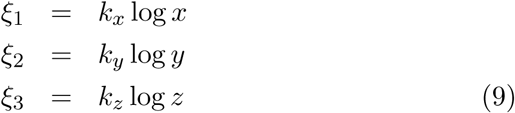

Which is just Fechner’s law applied to each coordinate.

There is nothing to guarantee that finding such a coordinate system is possible without adding extra coordinates. This is something that might be familiar from basic cartography, where a fundamental problem is that it is impossible to make a flat, two dimensional map that directly and simultaneously reproduces the geometric features of a globe (angles, area, distances etc). The question of whether it is possible to make a perceptually uniform colour space without adding coordinates has been tested empirically in the case of human colour vision, and various authors MacAdam [9], Wyszecki and Stiles [15] has shown that a uniform human chromatic plane cannot be represented in two dimensions. This does not mean, however, that a perceptually uniform space is impossible, only that it might need to be contained in a higher dimensional space. For example, the MacAdam [9] chromaticity space is a 2D surface which can be constructed in 3D space without any distortion (MacAdam [9] contains an image of a paper model that does just this, reproduced in Wyszecki and Stiles [15]).

## 3 Statistical Interpretation of Discrimination in Colour Spaces

We’re now in a position to look at the statistical foundation of these colour models. The general principle is that there are various sources of noise found in the environment and within the neural processes of the subject that limit the ability of the subject to fully distinguish between similar, but physically distinct colours. We can represent the information that the subject has available as a probability distribution over some ‘internal states’. Although at this point we do not need to specify exactly what these states are, it certainly wont hurt to think of them as the possible states of an ensemble of colour encoding neurons. Let’s call the internal state *X* and write its probability density as *p*(*x*; *ξ*). The parameter (coordinate) *ξ* denotes the particular colour that is being used as a stimulus.

We may also treat a behavioural outcomes the same way: as a random variable parameterised by the stimulus.

I have already outlined the significance of covariance matrices in current definitions of colour space. In the notation here the covariance matrix of *X* is given by.

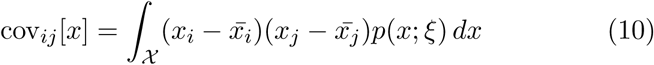

with *x*_*i*_ being the *i*^th^ observed component of *X* and 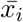 being its mean.

It might be clear to some from the outset that not just any covariance matrix of an internal state or behavioural response is up to the task of defining a colour space. The ‘units’ are all wrong! *ξ* is going to be a measure of the stimulus colour e.g. the number of photons absorbed by each type of photoreceptor per second, but *X* could be pretty much any physiological parameter, the potentials of an ensemble of neurons for example. There’s not even anything to guarantee that *X* has the same dimensionality as *ξ*.

Whilst the covariance matrix associated with an internal or behavioural state usually has the wrong units and dimensions, a covariance matrix that describes variation colour in colour matches does not have this problem; To get this kind of data one could present subjects with two colours, one which is fixed and another which they can modify, ask them to adjust another until it appears indistinguishable, and gather statistics on the distribution of the colours they choose; the results of this would necessarily be expressed in the right terms.

There are some parallels here with the process of statistical estimation. If we had direct access to the state *x* and wanted to find the value of *ξ* responsible for it, doing so would require us to construct some procedure to convert the *x* value we observe into an estimate of *ξ*. Mathematically, this procedure is just a function that takes in *x* values and gives us back estimated *ξ* values: an estimator in statistical parlance. In line with the conventions for estimators, I will denote this function 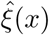 here, but we should bear in mind that in the case of the colour matching procedure above, the experimental subject performs the role of the estimator themselves and we can view the matches they make as estimated values, i.e. 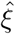.

Unlike the coordinates *ξ*, estimates 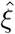 are functions of the random variable *x* and so are themselves random variables. This means that we can get various statistics from them, such as their mean and, most importantly, a covariance matrix. As we discussed above the inverse of covariance matrices describe an ellipse with a radius corresponding to the standard deviation, so it might seem like a good starting point for the metric tensor of a space. However, there is an issue with this from the geometric perspective: the inverse of the covariance matrix is not covariant; when calculate the covariance matrix in different coordinates, it changes in way that is in general only approximately contravariant (and thus its inverse is also only approximately covariant).

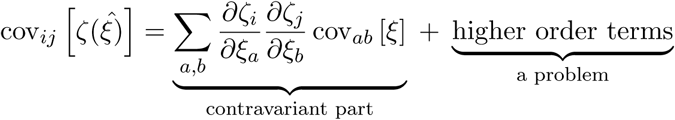

This is not only a mathematical technicality. The experimental consequence of this is that if one performs a matching experiment using one set of coordinates and then changes coordinates as one does in differential geometry, the results will not match up with an experiment performed in the second set of coordinates, that is, unless the higher order terms just happen to be zero (the conditions for which are outlined in Amari and Nagaoka [1]).

This said, at this point, an empirically minded reader might question the significance of this result. Perhaps the covariance is suitably small and the transformation sufficiently ‘forgiving’ for the higher order terms to be negligible, at least with the margin of experimental error. I have two things to say in response to this: Firstly, for the sake of the validity of existing results I hope this is indeed the case. Secondly, the magnitude of the error does not bear on whether it is mathematically correct, and we must be careful to distinguish the formal mathematical properties from their practical application. The fact is that there is a problem here, the only question is whether it is worth doing anything to remedy it. This is not a question I will dodge. After I have outlined a way of fixing this issue, I will present the case for its usefulness on grounds separate from the technical issues I have outlined so far.

My fix is suggested by the *Cramér-Rao bound* : a well-known result in statistical estimation theory which relates the covariance matrix of an estimator to a quantity called the Fisher Information, denoted here as ℐ.

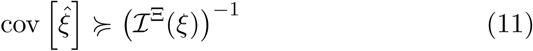

The Fisher information owes its name to R. A. Fisher, who defined it as the (co)variance of a quantity called the ‘score’, which is the derivative of the log probability with respect to the parameters. This leads to the definition:

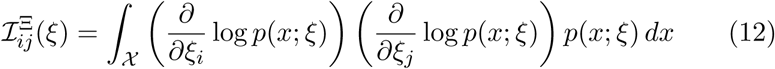

Unlike the covariance matrices of random variables we have discussed so far the Fisher information is not an observable variable: it cannot be *directly* accessed by behavioural experiments. This is because it is not based on the values taken by the random variable but on a the variable’s probability distribution, which itself can only be estimated. This makes it more suited to being calculated ‘from the bottom up’ based on physiological considerations (see section **??**) rather than from behavioural data.

Unlike the inverse covariance matrix of matches the Fisher information transforms as a metric tensor should:

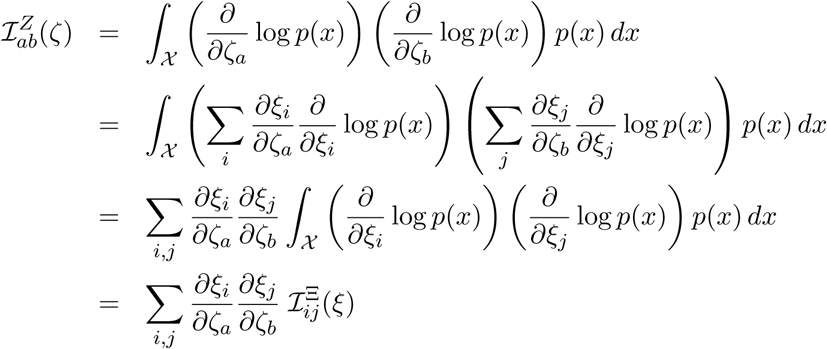

The Cramér-Rao bound states that Fisher information gives a lower bound to the covariance of an estimator, or equally, that if our subject is can never be more precise than the Fisher information allows whilst still being accurate. In terms of ellipsoids, this states that the ellipsoid representing the covariance matrix always contains the one representing the inverse Fisher information. The case of equality in the Cramér-Rao bound can be thought of as having chosen a coordinate system for representing the experimental stimulus in which can the informational limits of behaviour are properly captured by its variation. I will explore the details of this in the next section, and elaborate on what I mean by informational limits.

## 4 The Journey of Colour Information

Unlike covariance matrices for internal states (and behaviour in general) the Fisher information is always measured in terms of the parameters used to describe the stimulus and so one can use any set of physiological properties or behaviours to construct a colour space using the Fisher information. This is in contrast to a colour space based on covariance matrices where one has no choice other than to select a random variable which already has the same ‘units’ as the physical stimulus – in practice, this will be matching decisions or something deduced from discrimination behaviour.

This is of particular significance when we begin to think about the generalisability of experimentally determined colour spaces. Preferably, a colour space should not be merely a summary of experimental data in one particular situation, but something that applies in multiple situations, and ideally, in any situation one chooses. When we construct colour spaces we are aiming at something deeper than a summary of a experiment, we aspire to discover a common core, less narrow and superficial than, for example, the ability to distinguish two coloured semicircles. The advantage of using the Fisher information is that it gives us a way of talking about colour space at any point in the transduction process, right up to and including behaviour, so we could in principle apply it directly to that hypothetical point in the journey of colour information just before it branches into specialised task-specific processing.

Whilst the picture of a general process that splits into specialised processes at a single point is undoubtably simplistic, the idea that we can use the Fisher information to make a colour space at any point during processing of colour information is a powerful one. A general principle holds: we can look at the early processes and get a more general space, or look at the later ones for a more task-specific one. In fact, as I will now outline, there is a relatively straight-forward connection between the colour spaces at different stages of processing which can be used to think about this kind of problem; one which should be quite intuitive to those familiar with psychophysics.

### 4.1 Changing Spaces

Assume for the moment we can identify route through the brain from a colour stimulus ultimately leading to some observable behaviour, along which we can identify different states, each one determined by the previous one, with or without the addition of some amount of noise. Label these states *X*_1_, *X*_2_ etc. As an example: *X*_1_ might be the amount of light absorbed by photoreceptors, *X*_2_ might correspond to a population be bipolar or ganglion cells within the retina, *X*_3_ may be the optic nerves, with further stages processing through the visual cortex and the rest of the brain.^2^. This the natural way, I think, that most people would conceptualise the passage colour signals – or in fact any sensory signal – through the various stages of information processing.

The procedure I have just outlined is not a rigid formulation. There is some freedom in which states to include or not, how finely we wish split the path, and exactly which route we take. Additionally, it might be the case that some *X*_*i*_ could be chosen extended over time, or include extra parameters. This flexibility is important, as the history of colour vision has shown that it is in fact quite difficult to talk about a single, general purpose space. This said, it will almost certainly be the case that fully and accurately describing each *X*_*i*_ in detail for a real system will be quite challenging (though unnecessary), especially as one moves further into the brain; as such, in a practical application, the last *X*_*i*_ will likely be a generously filled black box. Such details, however, are not particularly relevant to the point at hand.

As long as we have a path constructed so that each stage (*X*_*i*_) is determined by a combination of the previous stage (*X*_*i*−1_) and noise we can write an inequality (in the sense of section 2.1.1) for the relevant Fisher information. This can be obtained from some basic properties of the Fisher information (see footnote^3^)

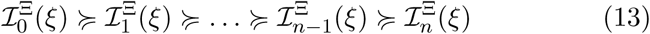

where 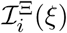 is the Fisher information at point *ξ* based on stage *X*_*i*_. This shows that as one moves along the path from stimulus to action the ellipses of fixed discriminability are growing (or more precisely, not shrinking) relative to the parameters – each one containing the previous. Conversely, the corresponding uniform space is always shrinking.

At the stimulus the ellipses are contracted to points, and every colour is perfectly discriminable from every other. The corresponding points in the uniform space are infinitely far from every other one. Figure 4.1 illustrates how the ellipses grow and the uniform colour space deforms and shrinks as we pass through the different *X*_*i*_s.

**Figure 3:**
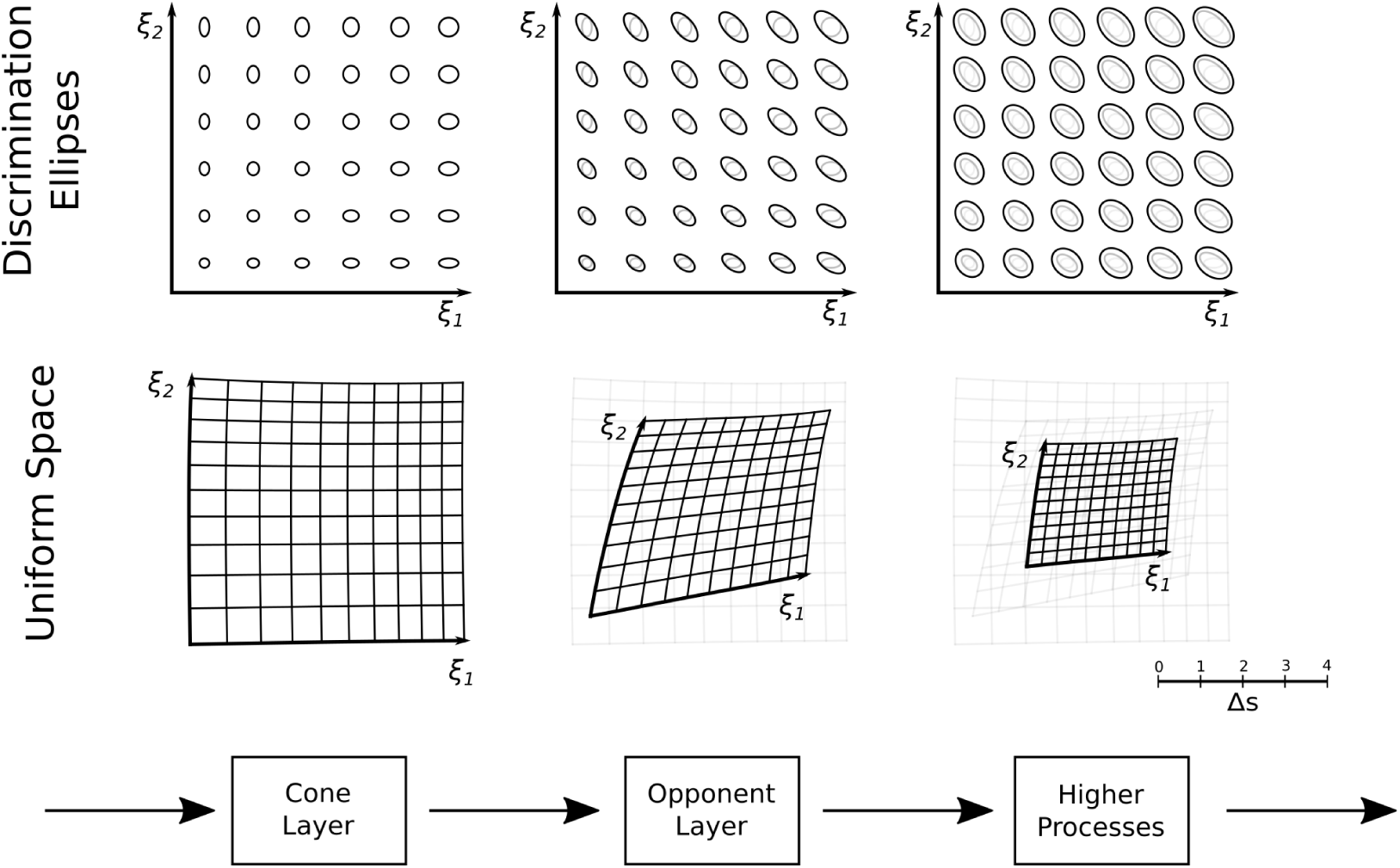
Illustration of the evolution of a colour space. The discrimination ellipses at each stage contain the ellipses of the previous stage, and the corresponding uniform colour space shrinks relative to the previous one.

**Figure 4:**
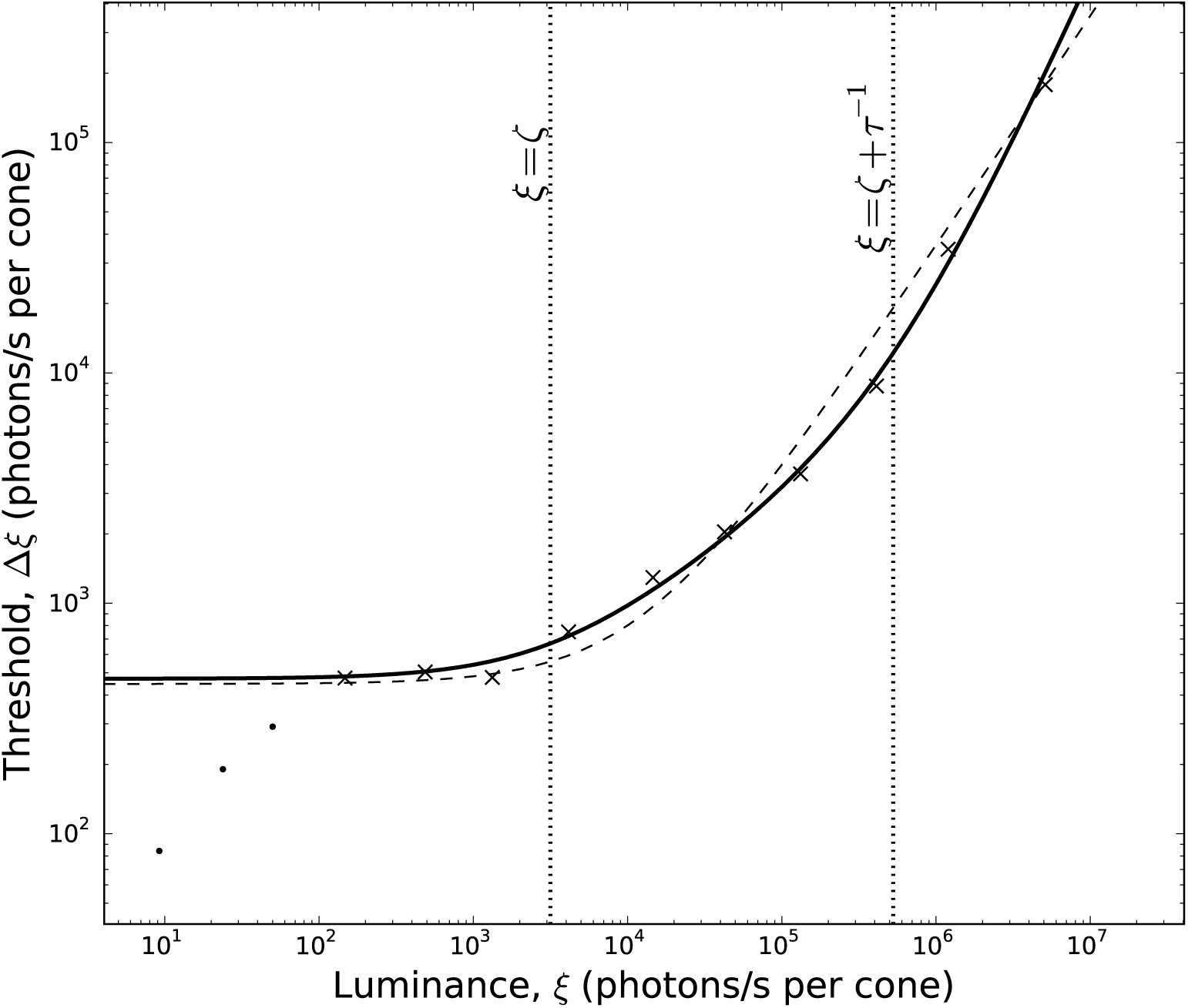
Stiles’ human contrast sensitivity data [4] with the model in section 6 (solid line) and Weber’s law (dotted line). Both fits are good but our model is better. The experiment determines the amount of 580nm light needed to discriminate from a 500nm background. At 500nm the long and medium cone excitations differ by approximately 0.3 log units so the degree of smoothing due to different excitations is very small. We may safely assume that short wavelength sensitive cones, being far less populous and less deferentially excited at 500nm, make very little contribution to the shape of the sensitivity curve. The data has been split into rod (dots) and cone (crosses) components for fitting. The constants used are, *τ* =1.9 × 10^−6^ s cone and *ζ* =2.3 × 10^3^ s^−1^ cone^−1^. The constant of proportionality is 1.7 × 10^−4^. The value of *τ* is significantly different from that associated with reported recovery half-lives, however, additional calculations based on model which includes deactivation and recovery rates separately gives more favourable values.

Returning now to the ideas of optimality and generality. Experiments that give information about a particular stage are ones where the inequalities to the right of that state in equation 13 are, in effect, equalities.

With regard to the idea of a general purpose space, we can think about what might be achieved using behavioural/psychophysical techniques. The inequalities above give some insight into how we might move in this direction, and the difficulties (and perhaps futility) of such attempts.

If we consider multiple behaviours, we would have multiple paths, but to some degree we would expect the earlier stages to be consistent. Given a set of different measured ellipses we can use the inequalities above to postulate a stage with ellipses are as large as possible without going outside the measured ellipses – whether this would match up with physiology is a different matter.

Although I have presented the case a single linear path in order to show some of the nicer properties of the Fisher information, it is certainly possible to think about more complex networks of information in similar terms. An example of case where such a network might be might be necessary would be in describing the disappearance of categorical boundaries between colours during ‘interference’ tasks[**?**], where it seems one would need at least both a linguistic and a non-linguistic pathway.

## 5 Covariance Ellipses and Fisher Ellipses

Ellipses based on the Fisher information differ from those that quantify empirical variation in matches or obtained from discrimination experiments. This difference is perhaps not obvious in the case of well designed experiments, but can be quite significant when this is not the case.

One way of thinking about the difference is that the Fisher ellipses describe the set of parameters that are confused, whereas the others describe what they are confused *as*. Hopefully this will become clearer with the following example.

Consider a matching experiment like described above, and two kinds of ellipses: one kind based on the variation in matches, and the other based on the Fisher information. In a ‘good’ psychophysical experiment these two should be comparable and the matching variation should reflect the confusion of stimuli. But now, let’s imagine something that would ruin any serious psychophysical experiment: a very persuasive experimenter enters the room and tells the subject that the experiment has been cunningly contrived, and although the colours of the set light might appear to be different to them, they are in fact white. In response to being told this, the subject abandons their original strategy and instead sets their light to exactly the same white on every trial.

Setting aside the plausibility of this situation, we can think about what happens to the two kind of ellipses. The ellipse describing the variation of the match becomes centred on white and at the same time shrinks to a point^4^: high precision, but low accuracy. The Fisher ellipse has the opposite behaviour: as it describes the parameters that are confused in the experiment and every colour is judged to be equivalently white, it grows to encompass all of the colours used in the experiment.

So what happened here in terms of information? It would be hard to argue that the experimenter did not give our subject any information: the effect was to increase the precision of their estimate after all. Perhaps we could instead say that it wasn’t information, but “false information”? This might be a better description, but what does ‘false’ mean in this context? It would be better to say that the experimenter gave the subject *information that was not dependent on the stimulus*. It seems that the Fisher information captures this falseness better than the estimate ellipse. If we think about the information flow from the section above, we can see that what has happened is that all the information about the stimulus has been thrown away by the time we get to the matching behaviour, the ellipses have grown huge and the colour space has shrunk to a point.

The example I have given is quite extreme, but we can imagine that with regard to any particular stage in processing, any departure from equality in equation 13 for the later stages, or any choice of parameter that fails to achieve the Cramér-Rao bound will result in this kind of effect to some degree.

## 6 Application

To illustrate how one might use the ideas I have outlined in practice I will derive a uniform colour space based on the first stage of phototransduction for which the uniform coordinates have a simple form. Models of this kind should be particularly useful for those in animal colour vision where physiological data is more easily obtainable than psychophysical. Despite its simplicity, this model will turn out to be quite good at predicting certain psychophysical results.

As this model only uses the first stage of phototransduction and no combination of colour channels has yet occurred, the channels can be analysed independently of each other. As the stochastic variation in channels is not correlated the task at hand reduces to finding the Fisher information for a single chromatic channel and the the Fisher information is a diagonal matrix.

Consider a fixed sized population of opsin molecules which can be either inactive (*X*) or active (*X*\*). In this population *X* is transformed to *X*\* at a rate proportional to the amount of light *ξ* plus a degree of spontaneous activity *ζ*, and each active molecule has a time constant of *τ* (half-life of *τ* log 2).

This process can be written as a chemical equilibrium with a forwards rate of *ξ* + *ζ* and a backward rate of 1*/τ*:

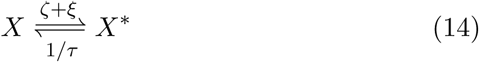

Let us assume that when viewing our stimulus an equilibrium is reached and that statistical fluctuations the amount of active opsin molecules (*X*\*) is the major source of relevant to discrimination. This assumption allows a time independent solution appropriate for making a colour space. This kind of model has been used before but not in the framework described here. Howard et al. [6] use it to measure the signal to noise ratio at the level of photoreceptor potential, by propagating the variance of the information limiting stage *‘forwards’* onto the membrane potential. Whilst their approach is appropriate for the kind of questions they ask, here we are do the opposite, taking the model of statistical variation *‘backwards’* onto the physical parameter to construct a colour space.

It can be demonstrated that in an equilibrium between two chemical species the numbers of the constituent molecules are described by a binomial distribution. The Fisher information for a binomial distribution is well known, but we must be cautious using reported values as the Fisher information is dependent on the choice of coordinate system. We have to be careful to keep track of this. The usual choice of parameter is the probability of the individual events, the *p* parameter. Superscript *P* to denotes this:

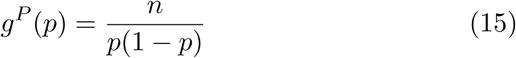

However, the parameter we are interested in isn’t *p*, but the rate of light induced photoisomerisation, *ξ*. We can rectify this can use the relationship between *ξ* and *p* to perform a tensor coordinate transform from the *p* coordinate to the *ξ* coordinate, i.e.

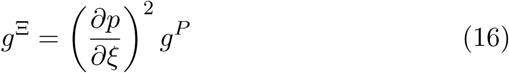

How are *p* and *ξ* related? We can find this out from the mass action kinetics of the species in equilibrium, where the rate of the reaction *X* → *X*\* is balanced by the reverse reaction *X*\* → *X*. This gives us an equation relating the amounts of the two species, written here as [*X*] and [*X*\*]:

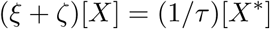

or equivalently

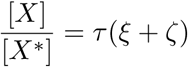

In standard chemistry terms *τ* (*ξ* + *ζ*) is the equilibrium constant and in the statistics of queues this quantity is called the “offered traffic”. It will be useful to just give this a single letter *φ*. To write this in terms of the parameters of a binomial distribution we can observe that in mass action kinetics the amount of active molecules [*X*\*] is the mean *np* and [*X*] is the what remains from the total, i.e. *n* − *np*, giving us:

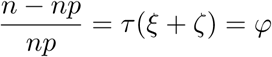

which, with some rearrangement, can be written as

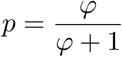

We can substitute this into equation 15 gives us the metric for the *p* coordinates written in terms of *φ*:

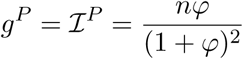

We can also use it to get the value of the derivative in equation 16:

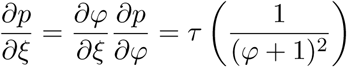

Putting all this together gives us a value for the Fisher information in the *ξ* coordinates:

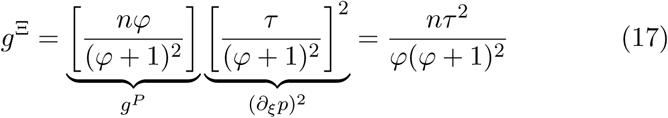

This expression can be understood as having three different “noise regimes, which can be better observed if we (1) look at the relative sensitivity, i.e. the ratio of perceptual difference to intensity difference, and (2) express it in terms of *ξ*.

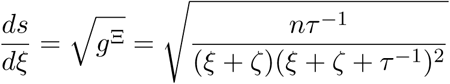

The first regime is when *ξ* is much smaller than *ζ* and this whole expression is approximately constant, this can be thought of as the ‘dark noise’ regime. The second regime when *ξ* is between *ζ* and *ζ* + *τ* ^−^1; This is where there is a square root relation between discriminability and intensity, it is the photon shot noise dominated region and corresponds to the Rose-de Vries law. Finally, for *ξ* above *ζ* + *τ* ^−1^ we have a three-halves power law. We can also see that this expression for the sensitivity increases with the square root of the number of molecules available.

We can also get a transformation into the corresponding perceptually uniform coordinates by integrating the corresponding line element

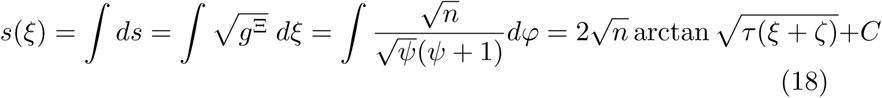

If we desire that *s*(0) is equal to 0 we can use the value of 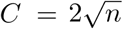 arctan *τζ*.

One feature of this model is that the available visual pigment gets less at higher light intensities resulting in decreased intensity. This is not a feature widely included in models of animal colour vision except for bees where there is a unique tradition regarding the quantification of colour[3]. Figures 4 and 5 show the result of this model for humans and bees respectively, and despite its simplicity it performs well in both cases.

**Figure 5:**
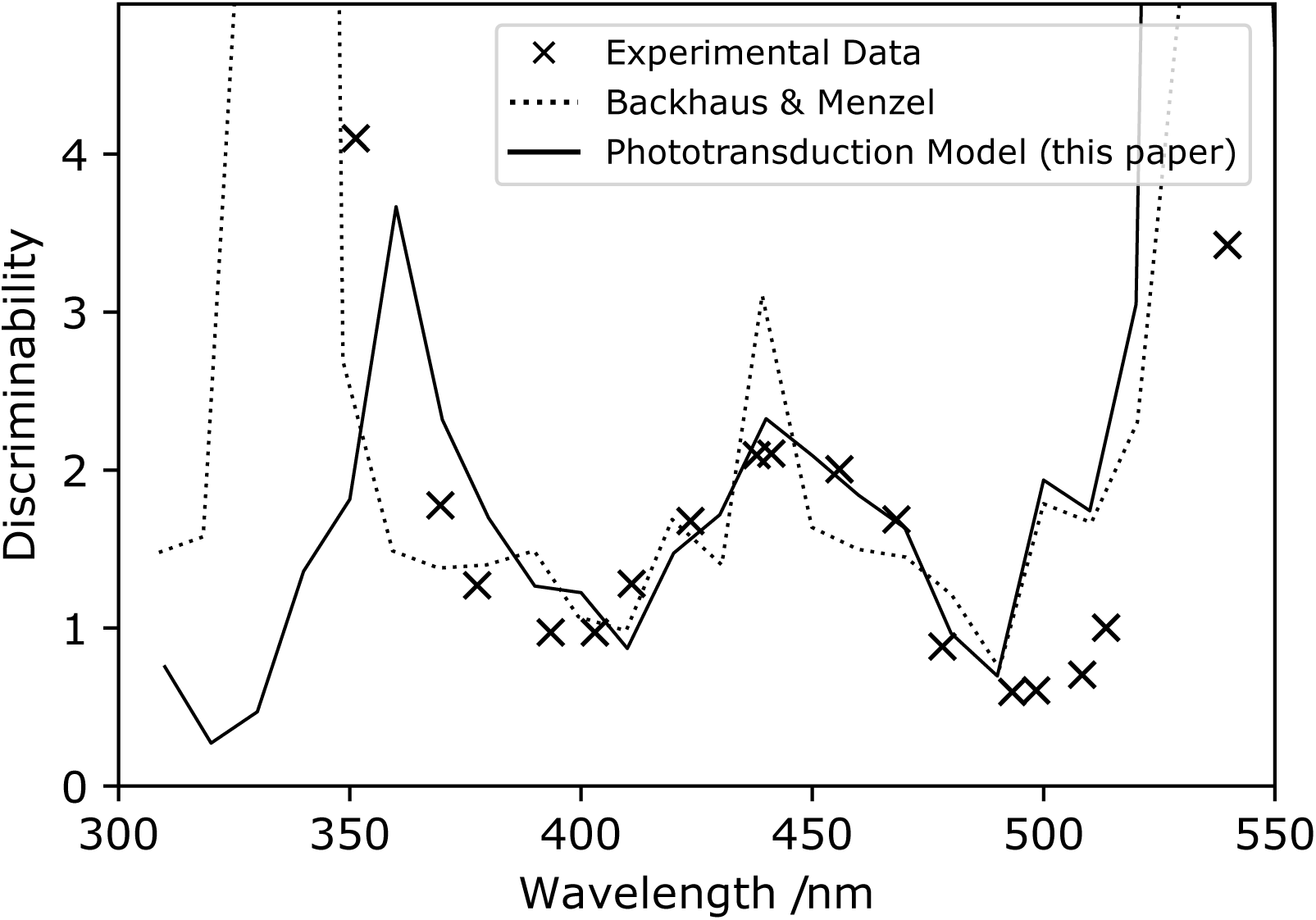
Application of the model from section 6 to wavelength discrimination (Δ*λ*) in honeybees (originally from [**?**]). This is based on the data reported in Backhaus et al. [2] and their model is shown here too. The model parameters were *ζ* = 0 and *τ* =5 s. Both the model in Backhaus et al. [2] and here are scaled to match the experimental data, and share common features that derive from the spectral sensitivity curves which they share[2]. Backhaus and Menzel’s model is based on neural excitation and noise and thus has similarities to the CIECAM spaces[10], and has been used to obtain the Honeybee Hexagon space which is widely used in bee vision[3].

It’s now worth revisiting the first point made in this paper and ask how these calculations would might have gone if we had tried to use the covariance matrix approach. This example makes the practical value of this approach quite clear. The variance of the amount of *X*\* is easy to calculate, and has a value often reported as *np*(1 − *p*). Similarly, it would be very easy to find the coordinate transform from *p* to *ξ* as we have above. It would be very tempting, then, to transform the reciprocal of the variance (1*/np*(1−*p*)) using the tensor transformation rules to get a metric in terms of *ξ*. But doing this yields a answer to qualitatively very different from the one arrived at above. It would be a factor of *n*^2^ smaller, the meaning of which is that the sensitivity decreases with the square root of the number of molecules molecules instead of increasing.

Clearly something has gone wrong here, and if we compare it to the Fisher information we can see what it was. The expression *np*(1− *p*) is only equal to inverse of the Fisher information when we use the right coordinates, and in this case the coordinate is *µ* – the mean amount of *X*\*. If we do the transformation from *µ* to *ξ* we get the same answer. But *without knowing the Fisher information we wouldn’t know which coordinate transform was the correct one to apply*.

This is the practical consequence of the variance being invariant with respect to the parameters as discussed above. With a covariance based approach, the coordinates are not directly ‘linked’ to the description of the stimulus, and without such a link one can only arrive at the correct answer by luck or good intuition.

## 7 Summary

The Fisher information provides an excellent way of grounding colour spaces in statistical theory. In this paper I have argued that conceptualising the colour spaces Fisher information solves mathematical issues with the usual formulation, as well giving us a more general and intuitive picture. I have also showed, by example, how it can help in practical applications.

Up to now I have mostly avoided talking about some of the wider implications of using the Fisher information there is a wealth of literature surrounding it and its relationship to statistics, information theory and decision theory. There are thus many ways this approach could be taken further. For instance, the Fisher information is found in Jeffreys’ prior used in Bayesian inference[**? ?**], suggesting that we might think of a perceptually uniform colour space for a particular visual stage as one in which no region is ‘preferred’ over another. This it might be particularly fruitful mental tool for those thinking about the ecology of colour, and how visual systems of colour vision evolve.

Is this enough to say that colour spaces are defined by the Fisher information? Does it matter? The idea of a colour space will no doubt continue to evolve as it has done in the past, but I hope that this approach can be of some value, both for guiding how we think about the particular class of colour spaces where discriminability is central, and for building models of perception in humans and other animals.

## Footnotes

1 Helmholtz’ final result is more complex than the one I present here, but the relative simplicity of this model makes it good for the purposes of demonstration.

2 Another way of saying this would be that the variables *X*_1_ to *X*_*n*_ would form a linear graph in Pearl’s graphical framework for causality[12]

3 Equation 13 follows from three well known[1, 7] properties of the Fisher information (a) the Fisher information is additive for independent variables (b) the Fisher information is zero if a variable does not depend on the parameters, and (c) summarising a variable only ever reduces the Fisher information (in the positive-definite sense). The incorporation of noise into a system *X* can be thought of introducing a statistically independent and parameter independent variable *Y* giving a combined variable (*X, Y*), and then summarising them both with some deterministic function giving *F* (*X, Y*). From (b) the Fisher information of *Y* is zero, and so by (a) the Fisher information of (*X, Y*) is the same as that of *X*. By (c) the Fisher information of *F* (*X, Y*) must be smaller than (*X, Y*) and thus *X*.

4 It would also be perfectly reasonable to calculate the covariance of the match assuming that the mean is the parameters describing the stimulus i.e. assuming that the estimation is unbiased. This would exhibit similar features, but in a way less simple way and so making it less appropriate as an example.

